# Stimulating music supports attention in listeners with attentional difficulties

**DOI:** 10.1101/2021.10.01.462777

**Authors:** Kevin JP Woods, Gonçalo Sampaio, Tedra James, Emily Przysinda, Adam Hewett, Andrea E Spencer, Benjamin Morillon, Psyche Loui

**Affiliations:** Brain.fm; Wesleyan University; Boston University; Aix-Marseille University; Northeastern University

## Abstract

Background music is widely used to sustain attention, but little is known about what musical properties aid attention. This may be due to inter-individual variability in neural responses to music. We test the hypothesis that music can sustain attention by affecting oscillations via acoustic amplitude modulation, differentially for those with varying levels of attentional difficulty. We first show that heavily-modulated music improves sustained attention for participants with more ADHD symptoms. FMRI showed this music elicited greater activity in attentional networks in this group only, and EEG showed greater stimulus-brain coupling for this group in response to the heavily-modulated music. Finally, we parametrically manipulated the depth and rate of amplitude modulations inserted in otherwise-identical music, and found that beta-range modulations helped more than other frequency ranges for participants with more ADHD symptoms. Results suggest the possibility of an oscillation-based neural mechanism for targeted music to support improved cognitive performance.

---

Music often has practical uses beyond aesthetic appeal^1^, and from mothers’ lullabies to laborers’ work-songs, the music we make to fill these roles reflects its function^2,3^. One possible use of music is to aid cognitive performance^4–6^. This has become increasingly important with the shift to knowledge-work^7–10^, along with widespread adoption of technologies like streaming and personal audio. To date, many different kinds of music have been used to aid focus in the workplace^11^. The diversity in music used for focus may reflect individual differences in cognitive styles: for example, personality differences are associated with the ability to sustain attention^12,13,14^. Preference and familiarity also contribute to effects of music on cognition^15–17^.

Another factor deserving special consideration is an individual’s ability to focus. Prior work has shown that auditory stimulation can aid performance in individuals with ADHD^18–21^. This has been explained by optimal stimulation theory, which poses that some individuals, specifically those with ADHD, require more stimulation than others to function best^22–25^. However, all these cases compare stimulation (music or noise) to silence, and no studies of this kind to date have used experimental conditions with different types of music.

If people who have difficulty focusing have distinct needs for focus-music, there is a chance to provide a targeted solution for those who could use it most. We were thus interested to see if music with different levels of arousal would affect people differently depending on their attentional capacity. If so, people with attentional deficits, such as symptoms of ADHD, may need specifically-designed focus-music. We hypothesize that arousal in music can affect performance differently in people with different levels of attentional difficulties^19,21,26^. Specifically, we hypothesize that more stimulating (i.e. more arousing) music should benefit people who experience more attentional difficulties as quantified by the ADHD Self-Report Scale (ASRS); i.e., high-ASRS individuals should improve in sustained-attention performance over time with more stimulating music.

We first compared performance (Experiment 1) on the Sustained Attention to Response Task (SART^27,28^) under three types of background acoustic conditions: AM+Music (i.e. music with fast amplitude modulations), Control-Music (with slow amplitude modulations), and Pink Noise. The AM+Music had fast modulations added that do not usually occur in music, and acoustic analyses (Figure 1) showed that despite similar frequency content, the tracks differed in the modulation domain due to this added modulation. We then used the same stimuli and task in experiments with fMRI (Experiment 2) and EEG (Experiment 3). Finally, additional behavioral experiments (Experiments 4A and 4B) tested the effects of modulation on sustained attention in an acoustically controlled manner.

**Fig. 1.**
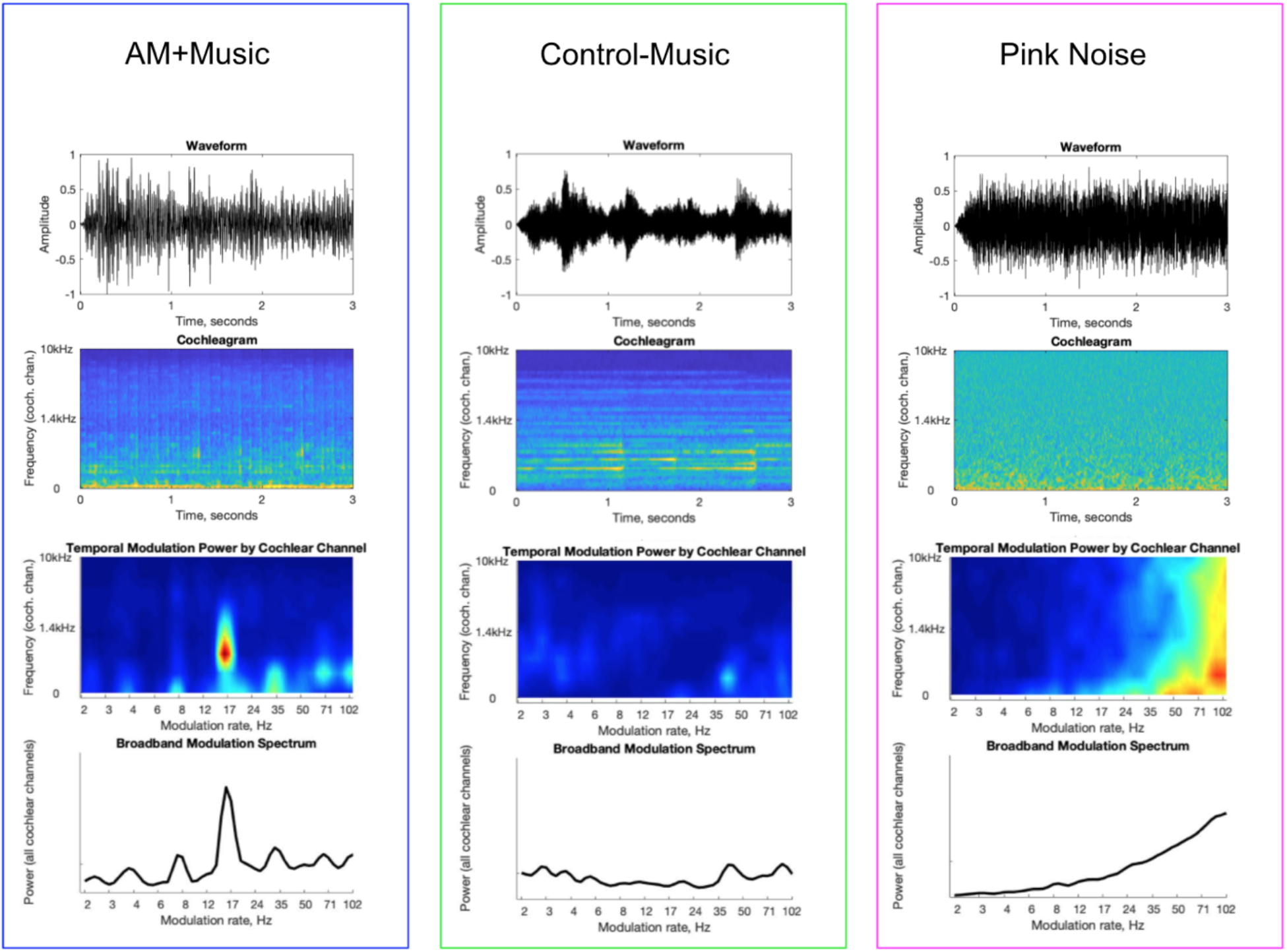
Acoustic analyses of auditory stimuli used in Experiments 1-3. Analysis of a 30-second excerpt from each stimulus type used in Experiments 1-3. Pressure over time (top row) first undergoes frequency decomposition via cochlear filtering. The energy in each cochlear channel varies over time (depicted on the cochleagram, 2nd row). These envelope fluctuations are then frequency-decomposed to produce a modulation spectrum representation (3rd row). The broadband modulation spectrum (bottom row) is the sum of modulation spectra across the cochlear channels. This broadband modulation shows peak in the AM+Music condition only, where rapid modulation was added to the music.

**Fig. 2.**
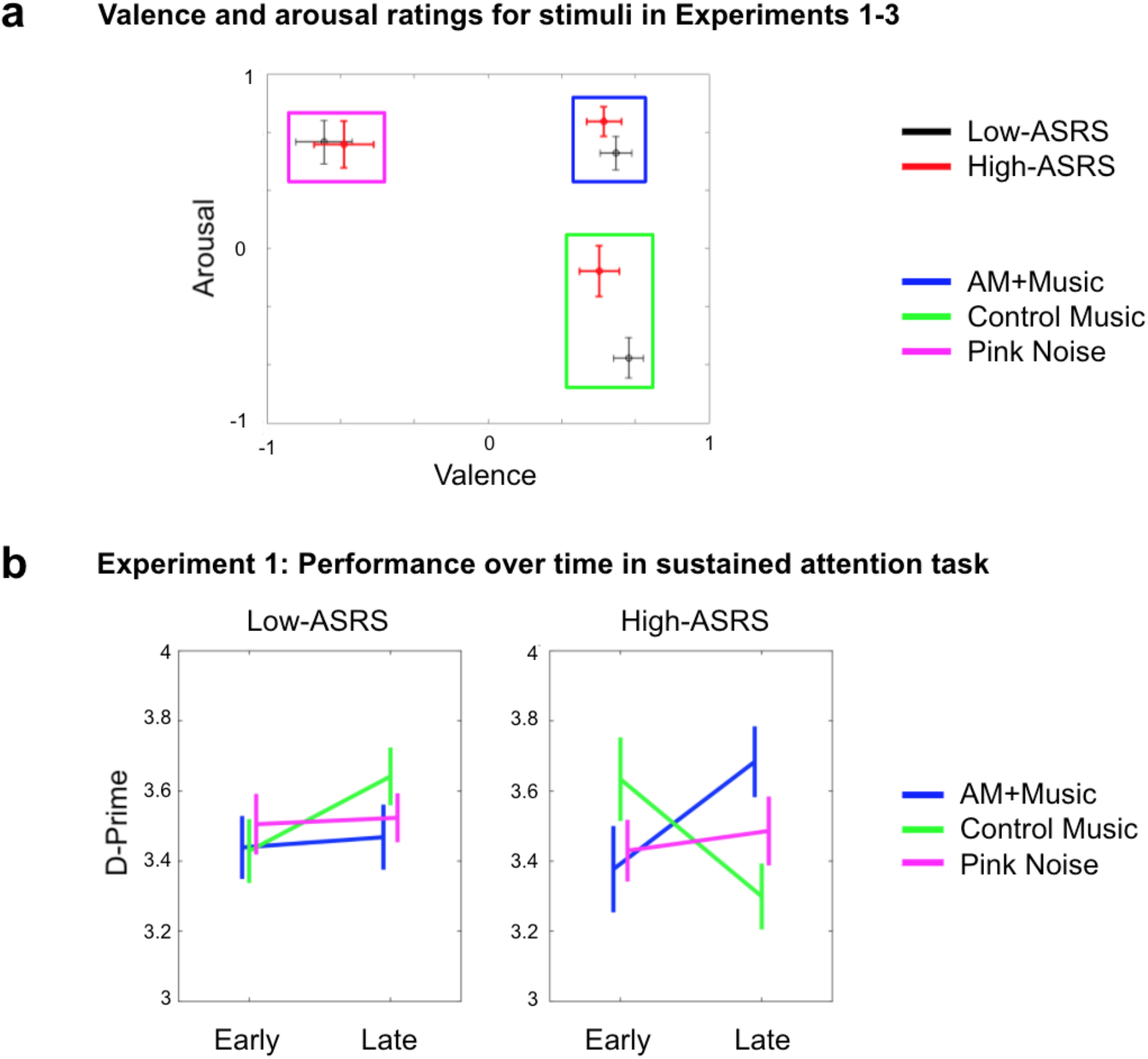
Stimulus valence and arousal ratings and performance over time in the Sustained Attention to Response Task (SART). **a**, Valence and Arousal ratings for the music used in Experiments 1-3, with listeners split by their level of self-reported attentional difficulty (ASRS score, median split). N=62 overall, N=31 in each group. **b**, Performance on the SART in a separate group of participants by music condition and by ASRS score. Each participant completed 3 blocks (the music conditions) presented in random order. N=87 overall, N=48 and N=39 in the Low- and High-ASRS groups respectively. Error bars depict +/- 1 standard error of the mean (within-subject).

## Results

### Experiment 1: Sustained Attention to Response Task (SART)

Participants were recruited and tested online via Amazon’s Mechanical Turk web service. The ASRS was obtained in all participants, and analyses were performed after a median split on ASRS score into high-ASRS and low-ASRS groups. In Experiment 1A, 62 participants (31 high-ASRS and 31 low-ASRS) rated the AM+Music, Control-Music, and Pink Noise for valence and arousal. In Experiment 1B, another 87 participants (39 high-ASRS and 48 low-ASRS) completed the SART under the three acoustic conditions.

Valence and arousal ratings for all three sound stimuli were entered into a multivariate two-factor mixed ANOVA, with the within-subjects factor of Music (3 levels: AM+Music, Control-Music, and Pink Noise) and the between-subjects factor of ASRS (high-ASRS vs. low-ASRS groups). A main effect of Music was observed on both valence and arousal (valence: F(2,120)=105,p<.001; arousal: F(2,120)=45,p<.001). A main effect of ASRS was significant for arousal (F(1,60)=6.0,p=.017) but not for valence (F(1,60)=0.166, n.s.). Participants rated AM+Music as positive in valence and high in arousal, Control-Music as positive in valence and low in arousal, and Pink Noise as low in valence but high in arousal. High-ASRS participants additionally rated the two music conditions (AM+ and Control-Music) as more stimulating (higher in arousal) than their low-ASRS counterparts.

Since the SART is a test of sustained attention over time, performance on the SART over time was analyzed with a 3-factor mixed ANOVA incorporating time as a within-subjects factor (2 levels: early vs. late trials), Music as a within-subjects factor (3 levels: AM+Music, Control-Music, and Pink Noise), and ASRS group as a between-subjects factor (2 levels: high-vs. low-ASRS). Overall performance (d-prime) in the two groups was not different (High-ASRS: mean = 3.50, stdv=0.71; Low-ASRS mean = 3.48, stdv=0.75). A three-way interaction was significant (F(2,170)=5.02, p=0.008): the music affected high-ASRS and low-ASRS participants differently over time, with high-ASRS participants showing an improvement in d’ over time during AM+Music but not during Control-Music. In contrast, low-ASRS participants showed an improvement over time during Control-Music but not AM+Music, despite no main effect of ASRS group (F(1,85)=0.01, p=0.92) and no main effect of music (F(2,170)=0.02, p=0.98). Since these two stimuli were rated as different in arousal but not in valence by both groups, differences in arousal may be related to these different patterns of SART performance over time.

The direction of this interaction--with fast modulations benefiting high-ASRS listeners in particular--aligns with our hypothesis that arousal in music can affect sustained attention differently in people because the optimal level of stimulation is greater for those with attentional deficits. To better understand the neural bases of this effect and how this might differ across the ASRS groups, we ran the same task and background conditions in neuroimaging experiments, with fMRI (Experiment 2) and EEG (Experiment 3), looking not only at the brain’s response to the different types of music, but also task-related activity.

### Experiment 2: SART fMRI During Background Music

In an fMRI study, 34 participants completed the SART under the same three background music conditions used in Experiment 1: AM+Music, Control-Music, and Pink Noise.

A within-subjects ANOVA comparing overall brain activity during the three conditions showed significantly higher activation during the AM+Music condition than in the other two conditions (p<.05 FDR-corrected) in multiple regions including the bilateral superior temporal lobes, frontal lobes, parietal lobes, and mesial and lateral occipital cortices, encompassing the default mode, executive function, and salience networks (Figure 3A). No other contrast showed positive suprathreshold clusters (all p>.05 FDR-corrected).

**Fig. 3.**
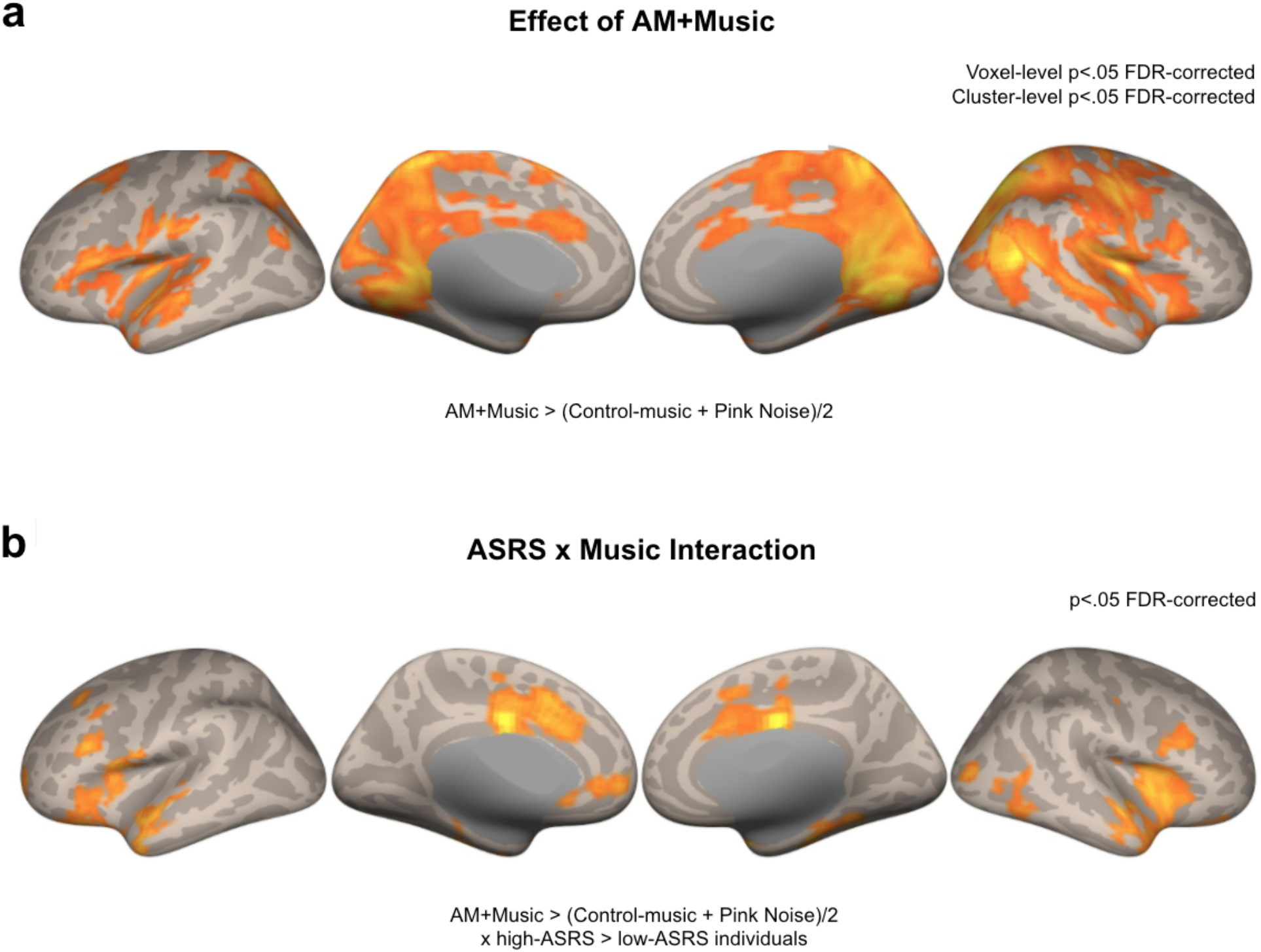
fMRI results comparing AM+Music, Control-Music, and Pink noise during the Sustained Attention to Response Task (SART). **a**, Contrast between AM+Music and the average of Control-Music and Pink Noise while participants performed a sustained attention (SART) task in Experiment 2. Higher levels of activity for AM+Music are widespread across many regions. **b**, Interaction between group (high-minus low-ASRS) and Music (AM+Music minus other conditions) was significant in the midcingulate (sometimes referred to as dorsal anterior cingulate; implicated in reward-mediated responding), anterior insula and ventral parietal areas (classic regions of the salience network), and middle frontal regions and anterior temporal poles. All results are significant at the p < .05 FDR cluster-corrected level.

### Individual differences in self-reported attention difficulties

When we divided the participants by ASRS (median split, as in Experiment 1) and compared their main effects of AM+Music (AM+Music > (Control-Music + Pink Noise) contrast), both participant groups showed activity in occipital and medial parietal regions. The interaction between music and group showed higher activity in the high-ASRS group than in the low-ASRS group during AM+Music compared to the other listening conditions. This interaction was significant in the insula and the anterior cingulate cortex, regions in the salience network^29^, with high-ASRS group showing larger differences than low-ASRS group in these regions (Figure 3B). Additional areas that survived cluster-wise FDR correction included the bilateral middle frontal gyri and frontal operculum, medial prefrontal cortex, bilateral temporal lobes, and lateral occipital cortex. The latter regions are part of the default network and the ventral attention network^30^, and their additional involvement is consistent with the role of the salience network in facilitating attention resources and accessing the motor system upon the detection of salient events.

### Higher activity in motor network linked to successful behavior during AM+Music

To relate behavior to brain activity during the different music tracks, we fit separate parametric models in SPM12^31^ for hit trials and false alarm trials for each auditory condition. Brain activity during successful responses, quantified as a contrast between activity during hits and activity during false alarms, showed significantly higher activity during the hits in all auditory conditions (**Figure 4**). Importantly, the Hits vs. FA contrast at the p < .05 FDR-corrected level showed more significant clusters during AM+Music than during any other condition (Control-Music, Pink Noise). These clusters centered around the sensorimotor network (supplementary motor area (SMA), precentral gyrus (PCG), the salience network (anterior cingulate cortex, anterior insula), and the visual association network (lateral and mesial occipital regions). These differences were observed despite overall similar hit and FA rates across the auditory conditions (one-way ANOVAs all Fs < 1). While the interaction between auditory conditions and groups was not significant at the whole-brain level, both high-ASRS and low-ASRS groups showed more activity for AM+Music than for the other auditory conditions during Hit trials compared to FA trials (Supplementary Materials). Together, these results show that AM+Music is linked to higher levels of brain activity in multiple networks, especially during successful behavioral performance of a sustained attention task. Furthermore, differences in brain activity during AM+Music listening (under Hits>FAs) were observed between people who report high and low levels of attentional difficulties, with more activity observed in (which regions) in the high-ASRS group. These individual differences in brain activity could underlie group differences in performance from Experiment 1.

**Fig. 4.**
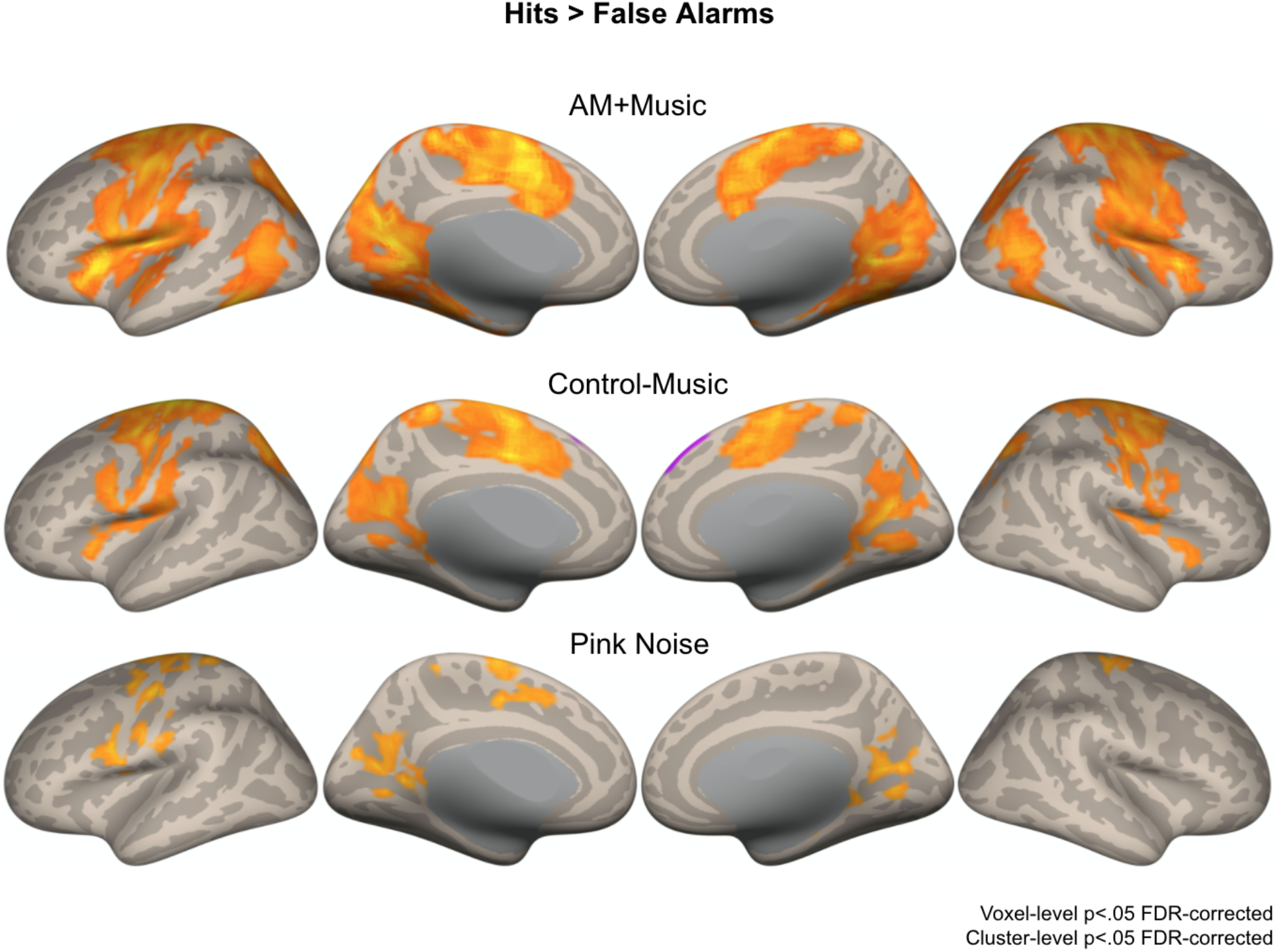
fMRI results comparing Hits and False Alarm trials from the Sustained Attention to Response Task (SART) during AM+Music, Control-Music, and Pink Noise. Brain activity during Hits contrasted against False Alarms on the SART showing increased activity during correct trials despite similar motor output, centering around the sensorimotor network and the salience network during AM+Music compared to Control-Music and Pink Noise. Warm colors show greater activity for Hits; cool colors show greater activity for False Alarms.

Since the musical stimuli were highly rhythmic, we expected that they might affect rhythmic activity in the brain, and that group differences in such activity could provide further insight into the mechanisms by which music affects sustained attention. To capture rhythmic neural activity and relate it to stimulus rhythms with high temporal precision, we turned to an EEG study with the same stimulus and task conditions as the fMRI study reported above.

### Experiment 3: SART EEG During Background Music

Forty participants had their EEG recorded while they performed the SART task under the same three background music conditions used in Experiments 1 and 2: AM+Music (containing fast modulation rates; more arousing), Control-Music (containing slow modulation rates; less arousing), and Pink Noise. Hit and FA rates were overall similar across the auditory conditions (one-way ANOVAs all Fs < 1).

### Stimulus-brain coupling EEG shows phase locking at peak frequencies of amplitude modulation

We assessed coupling between the EEG and the acoustic signal by computing the stimulus-brain phase-locking value (PLV) for every frequency (in 1-Hz bins). During the AM+Music, stimulus-brain PLV showed prominent peaks at 8, 12, 14, 16, 24, and 32 Hz (Figure 5A). Since the AM+Music was at 120bpm (i.e., quarter-notes at 2Hz), 8, 16, 24, and 32 Hz are harmonics of the note rate, previously observed to entrain cortical activity^32^, while 12, 14, and 16 Hz reflect the amplitude-modulated frequencies.

**Fig. 5.**
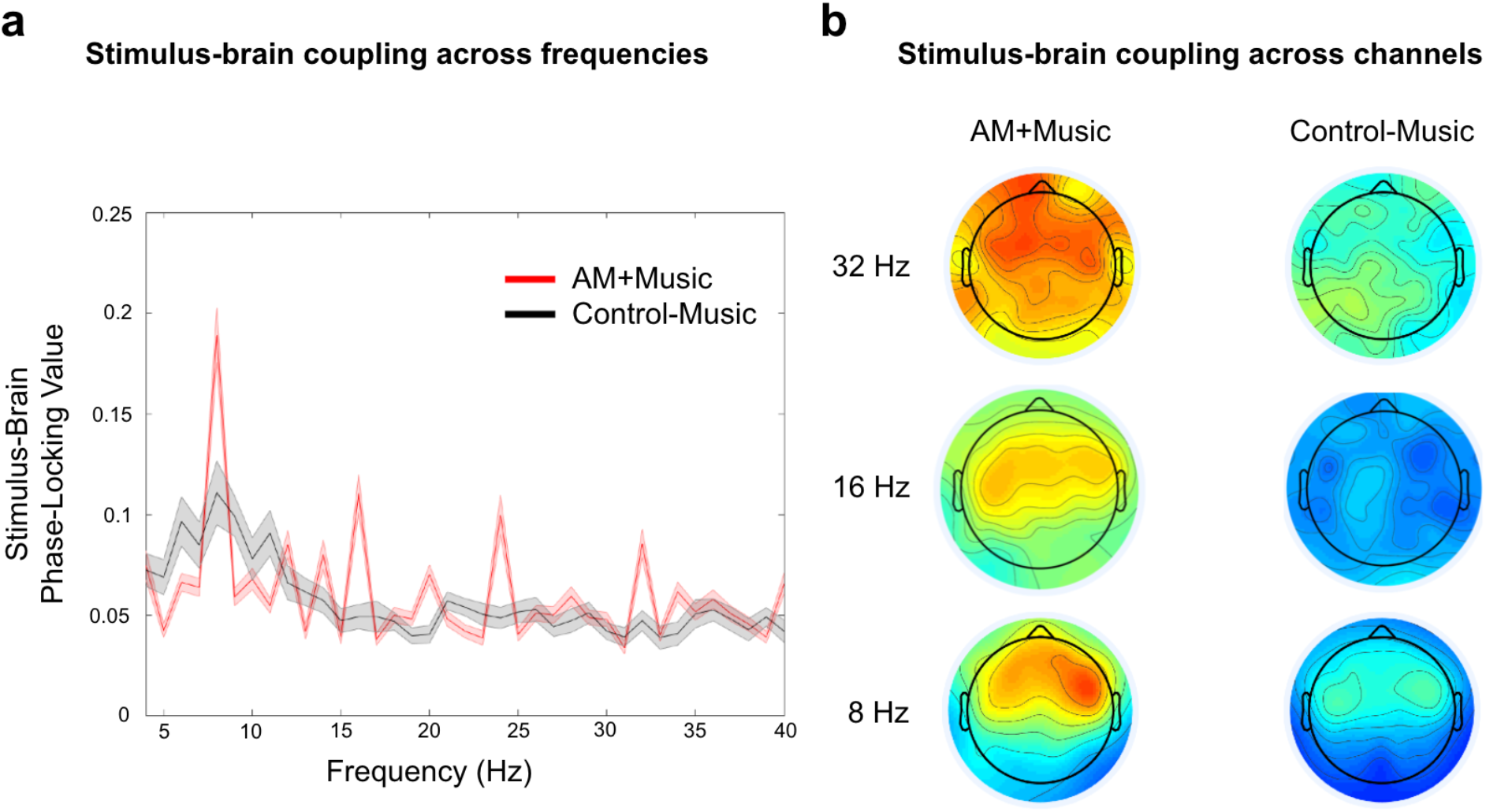
EEG reveals stimulus-brain coupling. **a**, For Experiment 3, we conducted hertz-by-hertz Morlet wavelet analysis of each acoustic stimulus (amplitude-modulated and control) and evaluated the phase-locking value (stimulus-brain coupling) between acoustics of the stimulus and the EEG. Red trace = Mean±SE Phase Locking Values (PLVs) of frontocentral electrodes for AM+Music. Black trace = Mean±SE PLVs of the same electrodes for the Control-Music. Results show peaks of phase-locking activity at the note rates and its harmonics (8, 16, 24, 32 Hz) as well as the amplitude-modulated frequencies (12, 14, 16 Hz). **b**, Topographic distributions of the peaks of phase-locked activity at 8, 16, and 32 Hz for amplitude-modulated and control stimuli.

PLV at 8 Hz during AM+Music was strongest at frontal recording sites. In contrast, PLV during Control-Music was much lower and less sharply tuned, reflecting less focused neural tracking of acoustic rhythms when listening to Control-Music. Looking across the whole brain (Figure 5B), PLV at 8Hz was stronger in the AM+Music condition, even over the same frontal recording sites. We computed an effect size (Cohen’s d) measure for every 4-Hz bin (thus capturing 8, 12, 16, 20, 24, 28, 32 Hz) across the full frequency spectrum, comparing AM+Music against Control-Music, averaging across all frontal, central, and parietal electrodes. This resulted in a Cohen’s d of 3.74, confirming a highly statistically significant difference between AM+Music and Control-Music at these frequencies of interest. In contrast, the same analysis for 1-Hz bins showed a Cohen’s d of 0.227, suggesting higher stimulus-brain coupling at multiples of 4-Hz, consistent with the note rate and with the amplitude-modulation rates in the music.

### Phase-locking across ASRS groups

We compared stimulus-brain PLV between high-ASRS and low-ASRS groups. The high-ASRS group showed higher PLV in high frequencies, especially during the AM+Music condition (Figure 6A). This pattern of higher PLV in high-ASRS participants was mainly observed during the second half of the AM+Music sessions (Figure 6c-d), suggesting that the effects of AM+Music emerge over time, consistent with the behavioral results from Experiment 1. This higher PLV over time for high-ASRS participants was observed at 12, 16, and 24Hz, and not observed below ∼12Hz, even though the 8Hz band in particular was strongly affected by modulated music in the overall PLV (Figure 6A), and lower modulation frequencies were still acoustically strong in the AM+Music (Figure 1A). Instead, the selective increase in phase-locking at higher modulation frequencies may hint at the role of particular oscillatory regimes in the behavioral effects of this music. Increments of 4 Hz Cohen’s d comparing AM+Music against Control-Music was 3.7391 for low-ASRS and 2.5161 for high-ASRS, but for increments of 1 Hz: Cohen’s d was 0.4587 for low-ASRS and 0.0296 for high-ASRS individuals. Comparing late vs. early in the listening session, Cohen’s d was −0.3960 for low-ASRS individuals, and 0.5343 for high-ASRS individuals when PLV was tested at increments of 4 Hz, but when PLV was tested at 1-Hz increments, Cohen’s d was - 0.1945 for low-ASRS and 0.0566 for high-ASRS individuals. The consistently higher effect size when PLV was tested at 4-Hz increments supports the finding that PLV was highest at integer multiples of 4 Hz. And while low-ASRS individuals were more effective at tracking the AM+Music than were high-ASRS individuals, the high-ASRS individuals showed more of a change over time, resulting in higher PLV late in the listening session.

**Fig. 6.**
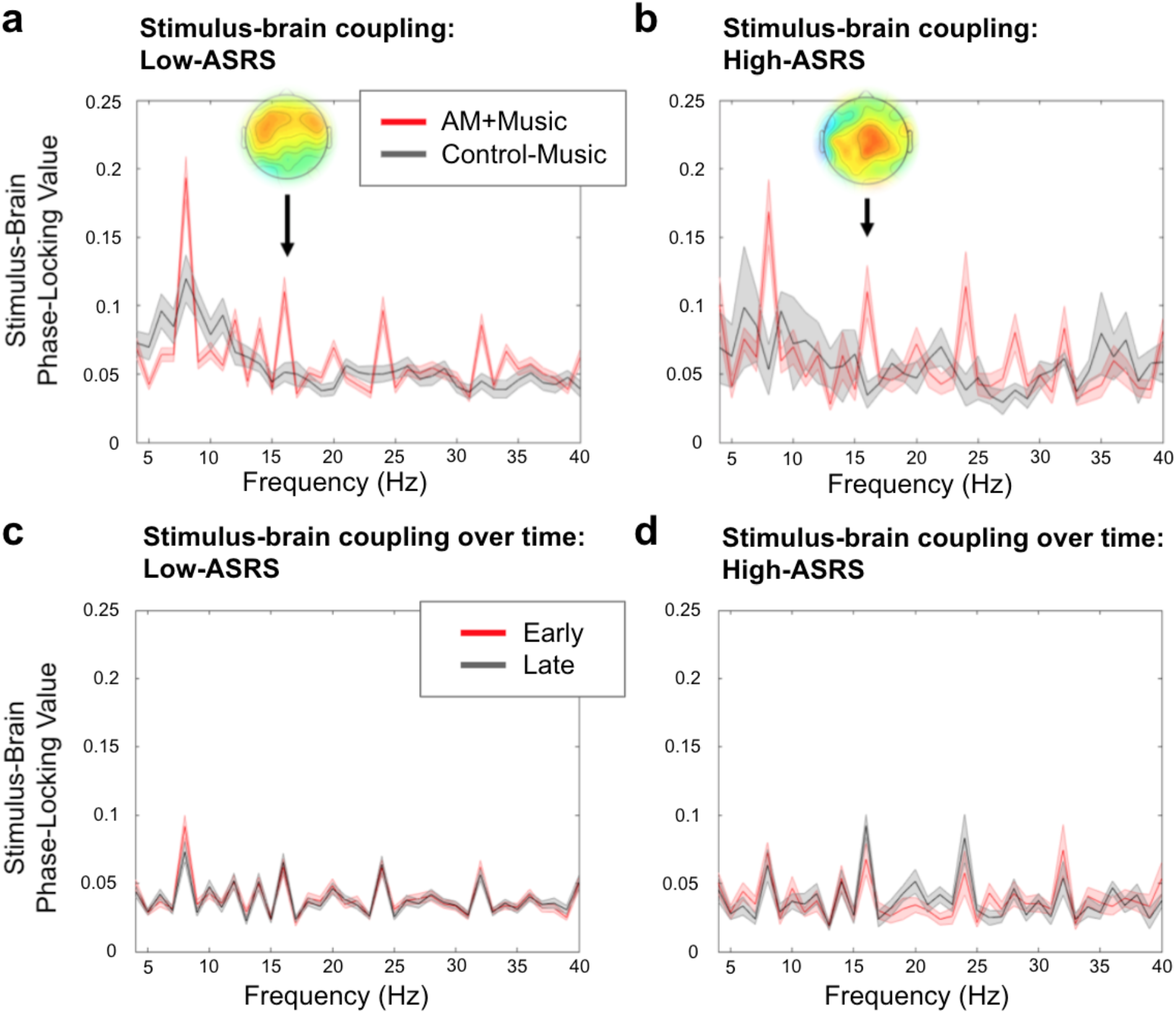
Stimulus-brain coupling for high-ASRS and low-ASRS groups. Top row: PLV during AM+Music (red) and Control-Music (black). **a**, low-ASRS participants; **b**, high-ASRS participants. High-ASRS participants show higher PLV in higher frequencies, especially while listening to AM+Music. The inset topos plots show the scalp distribution of PLV at 16 Hz during AM+Music. Bottom row: PLV during AM+Music listening in the first half (black) and second half (red) of the recording session. **c**, low-ASRS participants; **d**, high-ASRS participants. Black: first half of the session; Red: second half of the session. Higher PLV for high-ASRS participants emerge over time, and is higher late during the session. This is consistent with the behavioral improvement seen in the second half of the AM+Music session for high-ASRS participants.

### Experiment 4: Parametric manipulation of modulation rate and depth

The stimuli in Experiments 1-3 were taken from commercially available focus music that one might encounter if searching for music to work to. They were chosen to have a dramatic difference in modulation characteristics, as was apparent in acoustic analyses (Figure 1). However, they also differed in low-level acoustic properties (e.g., overall spectral balance) as well as musical features (e.g., tonality, instrumentation). To control for these differences and to isolate the effect of amplitude modulation, we developed new stimuli in which otherwise identical music was manipulated to impose modulation of varying rates and depths (Figure 7).

**Fig. 7.**
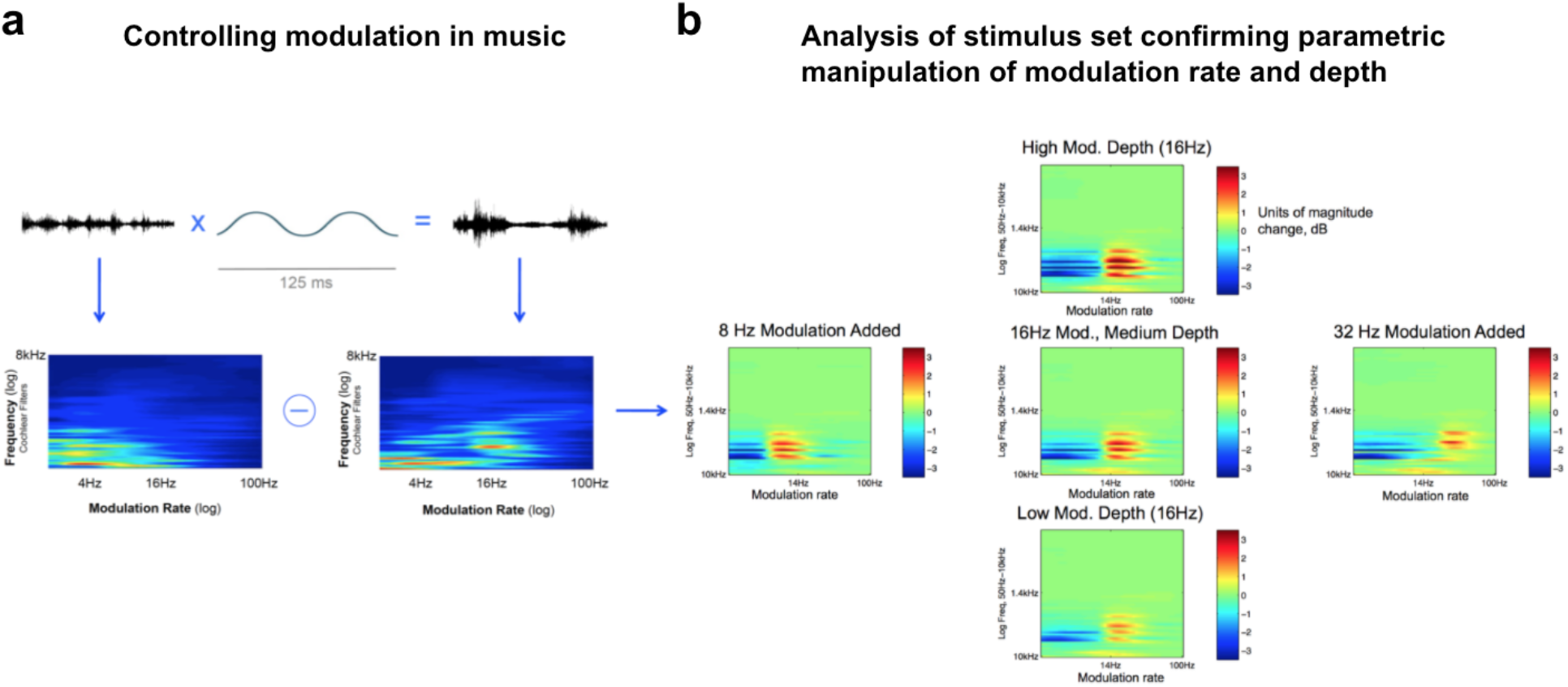
Controlling modulation in music stimuli. Modulation in sound can be summarized by the modulation spectrum: The vertical axis on these 2D plots depicts audio frequency (0-8kHz). The horizontal axis shows the amplitude modulation spectrum (amplitude fluctuations from 0-100Hz). **a**, Applying modulation to music results in a modulation spectrum with a peak corresponding to the rate of the added modulation. Upper: The pressure wave is multiplied by a modulator (in this case 16Hz) to produce a modulated signal. Lower: This pressure wave is bandpass filtered, and fluctuation rates in each channel are shown in the modulation spectrum. **b**, Validation of stimulus manipulations used in Experiment 4. The stimulus space is illustrated by these panels, each of which shows the difference in modulation spectrum (normalized power) between an unmodulated track and the experimental conditions derived from this track. Rate of added modulation increases moving rightward, while depth increases moving upward. The absence of differences elsewhere on the modulation spectrum shows that the experimental conditions were altered in a controlled way, with modulation properties different from the original only as specified (stimulus validation).

### Parametrically testing the effects of amplitude modulations: Rate and Depth

Acoustic amplitude modulations are known to drive neural oscillations, i.e., to induce a selective amplification of neural activity at this frequency. This effect occurs along the auditory pathway but also in cortical networks such as the attentional network ^33–36^, and may thereby impact cognitive processes. We chose to test rates of 8, 16, and 32Hz for two reasons: First, these fall within ranges of distinct neural oscillatory regimes that are known to have different functions in the brain. Alpha (8-12Hz), beta (14-25Hz) and gamma (25-100Hz) rhythms are three such ranges, and our experimental conditions using 8, 16, and 32Hz modulation thus fall into each of these oscillatory regimes. Second, these rates were chosen to correspond to note values. As we used music that was composed at 120 beats per minute (2 Hz), amplitude modulation rates at 8, 16, and 32 Hz correspond to 16^th^, 32^nd^, and 64^th^ notes respectively.

Beta-band cortical activity is implicated in the maintenance of sensorimotor or cognitive states^37^ and top-down processing in general^38^ including attentional control^39,40^. Moreover, in spatial attention tasks, beta-band increase is observed in the hemisphere that represents the attended stimulus, resulting in enhanced processing of the attended stimulus^41^. In contrast to the other bands, stimulating neural activity in the beta band with the 16Hz modulation condition thus seemed most likely to confer a performance benefit on sustained attention, and based on results from Experiments 1-2 we would expect to see a greater effect in our high-ASRS participants. We thus hypothesized that for higher-ASRS individuals, the 16Hz rate would produce effects on performance superior to a no-modulation control condition, while the other rates would not. We also hypothesized that greater modulation depth would have a greater effect on higher-ASRS listeners.

Depth of modulation refers to how heavily the sound is modulated, rather than the rate of modulation. A maximal depth of modulation would mean that sound energy is reduced to zero at the troughs of the applied modulating waveform, while a very low depth would mean barely-perceptible modulation. While a greater modulation depth is expected to impact neural oscillations more strongly, beyond a point the underlying music suffers aesthetically as the sound becomes distracting due to increased auditory salience from the sudden changes in loudness over time^42^.

175 participants (81 high-ASRS and 94 low-ASRS) completed two SART experiments online (4A and 4B) each with four conditions, testing modulation depth and modulation rate separately. In both cases two groups of participants each heard four conditions, presented for five minutes each in counterbalanced order. The first group of participants heard no-modulation, 8Hz medium-depth, 16Hz medium-depth, and 32Hz medium-depth modulation. The second group heard no-modulation, 16Hz low-depth, 16Hz medium-depth, and 16Hz high-depth modulation. Thus, rate and depth (4 conditions each) were tested on separate groups of participants, but each participant heard all four possible rates or all four possible depths for five minutes each. This limited the duration of each condition, but maximized statistical power in comparing across all the conditions in one dimension of the modulation spectrum, while controlling for intrinsic between-subject differences in performance that are unrelated to our conditions of interest.

Valence and arousal ratings for all six sound stimuli were entered into a multivariate two-factor mixed ANOVA (as in Experiment 1), with the within-subjects factor of Music (6 levels: No Modulation, 16Hz Modulation at Low, Medium, and High Depth, 8Hz and 32Hz at Medium Depth) and the between-subjects factor of ASRS (high-ASRS vs. low-ASRS groups). A main effect of Music was observed on both valence and arousal (valence: F(5,300)=32.3, p<.001; arousal: F(5,300)=7.1, p<.001). A main effect of ASRS was significant for arousal (F(1,60)=6.0,p=.017) but not for valence (F(1,60)=0.166, n.s.).

Overall performance (d-prime) for the two groups in the rate experiment: Low-ASRS, mean=3.68, stdv=0.72; High-ASRS, mean=3.36, stdv=0.69; and in the depth experiment: Low-ASRS, mean=3.56, stdv=0.52; High-ASRS, mean=3.52, stdv=0.84.

The modulation depth experiment (Figure 8B) yielded overall d-prime values of Low-ASRS mean=3.56 stdv=0.52; High-ASRS mean=3.52 stdv=0.84. The depth experiment revealed a trend toward an interaction between ASRS group and SART performance over time (3-factor mixed ANOVA; Between-Subjects factor of ASRS group: F(1,270)=3.70, p=0.070), but no effects of music condition. The modulation rate experiment (Figure 8C) showed a significant main effect of ASRS group (3-factor mixed ANOVA; between-subjects factor of ASRS group: F(1,80) = 4.14, p=0.045), but also found interactions between ASRS and Time (F(1,270) = 4.43, p=0.038), as well as ASRS and Music (F(3,240) = 2.89, p=0.036). This effect of Music by ASRS appears to be driven by high-ASRS participants’ better performance when hearing 16Hz-modulated music; this finding is consistent with Experiment 1 and suggests rate-specific effects of modulation in music, possibly subserved by oscillatory mechanisms in the brain.

**Figure 8.**
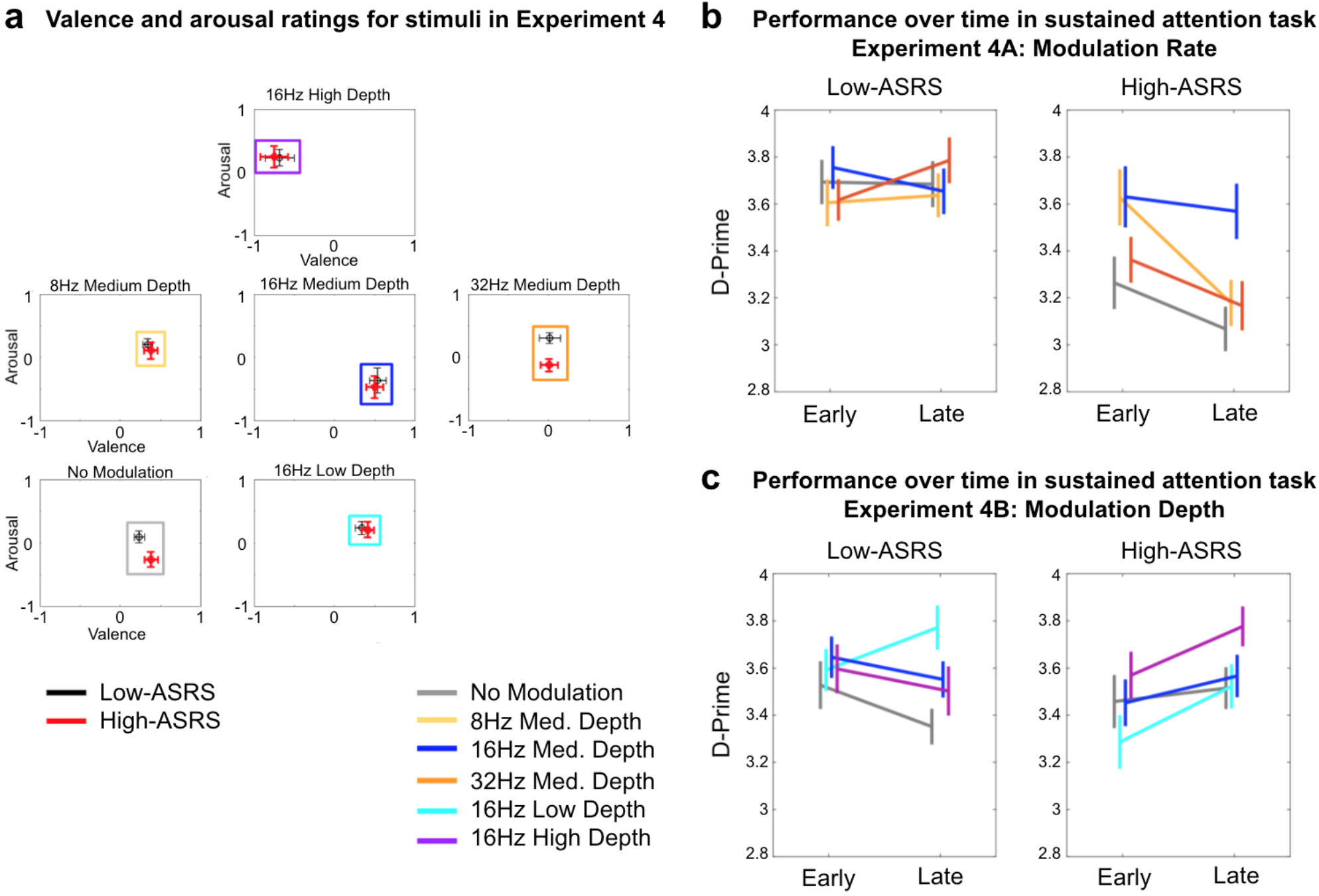
Stimulus valence and arousal ratings and performance over time in the Sustained Attention to Response Task (SART) for stimuli parametrically manipulated in modulation rate and depth. **a**, Valence and Arousal ratings for the music used in Experiment 4, with listeners split by their level of self-reported attentional difficulty (ASRS score, median split). N=62 overall, N=31 in each group (same participants as Experiment 1). **b**, Performance on the SART for varying rates of added modulation. N=82 overall, N=46 and N=36 in the Low and High ASRS groups respectively. **c**, Performance on the SART for varying depths of added modulation. N=93 overall, N=48 and N=45 in the Low and High ASRS groups respectively. In Experiments 4A & 4B, each participant completed 4 blocks (the music conditions) presented in random order; the blocks are overlaid in the figures.

## Discussion

Performance on a variety of everyday tasks requires sustained attention. Cognitive failures, specifically failures in sustained attention, are linked to mind wandering, which is associated with decreased productivity and happiness^43^. Music is widely used to help with sustained attention, with many use cases in everyday life ranging from café music to noise generators for work environments. Here, we show that effects of music on sustained attention depend on one’s level of attentional difficulty. In high-ASRS participants, who experience more attentional difficulties, a specific type of amplitude-modulated music (AM+Music) was more effective at engaging the salience, executive control, sensorimotor, and visual networks, and at coupling with rhythmic brain activity at multiple frequencies. While previous studies have suggested that differences in perceived valence and/or arousal of music may explain differences in sustained attention, our results show that valence and arousal do not explain effects of modulation across groups. Rather, the effect of music on sustained attention appears to result from specific interactions between functional differences in brain activity and amplitude modulation patterns in the musical sounds. Individuals who self-report attentional difficulties are more sensitive to the effects of AM+Music in behavioral and neural measures. Individual differences in sensitivity to background music may be attributable to “optimal stimulation level”^44,45^, i.e., the effect of external stimulation on cognitive performance, which differs across individuals, e.g. between introverts and extroverts^12,46,47^ and between groups with and without attentional deficits^25,26,48^. These stimulus-brain interactions could partly explain—in addition to preference and familiarity^15–17^—why people use such different types of music to help them focus, and may suggest routes to more effective personalized focus music in the future.

Behavioral experiments (Experiment 1) showed that high-ASRS participants performed better on a sustained attention task when listening to music that contains rapid amplitude modulations, compared to music with slower amplitude modulation or pink noise (broadband/noisy modulation). While the AM+Music also elicited higher arousal ratings than the Control-Music especially in the high-ASRS individuals, in a follow-up experiment (Experiment 4) high-ASRS participants did best in a condition with relatively low arousal; together the experiments suggest that the parameters of amplitude modulation causally affected behavior independent of arousal ratings. This suggests that music can be tailor-made with added acoustic modulations in order to aid performance in attentionally-demanding tasks, and that this can be done on an individualized basis with ASRS as an important factor.

FMRI and EEG experiments (Experiments 2 & 3) identify some mechanisms behind these effects. The fMRI experiment (Experiment 2) showed higher activity overall in response to AM+Music. For the high-ASRS group in particular, AM+Music elicited higher activity in the anterior cingulate cortex and the insula, core nodes of the salience network. Directly contrasting Hits against False Alarms showed a greater extent of correct response-related activity during

AM+Music than other conditions, particularly in motor regions. This could indicate that behavioral advantage for AM+Music relates to activity in the motor system, i.e. that the AM+Music could be more effective at priming the motor system to respond correctly when receiving the task-relevant sensory signals from the sensory cortex.

Our EEG experiment (Experiment 3) showed that phase-locking activity was strongest during AM+Music and particularly stronger in the high-ASRS group, in whom the phase-locking activity also increased over time. These results suggested that amplitude modulation could underlie the difference in performance between ASRS groups observed in Experiment 1. To isolate the effects of amplitude modulation, a final behavioral experiment was conducted using music that differed only in the rate or depth of added modulation (Experiment 4). Here we found that 16Hz and high-depth amplitude modulation resulted in overall best performance in the high-ASRS group compared to the other conditions, resulting in performance for high-ASRS participants that was similar to the low-ASRS group.

### Effects of music depend on one’s level of attentional difficulty

A consistent result across our experiments was that effects of the music varied depending on an individual’s level of attentional deficit, as measured by self-report on the ASRS. While previous studies have observed that individuals with attentional deficits are differently affected by music versus silence^5,14,49^ or noise^12,20,21,50^, here we extend the findings to show that differences between pieces of music are sufficient to affect sustained attention. In our EEG experiment (Experiment 3) we found that the difference between AM+Music and Control-Music was greater for the high-ASRS participants (Figure 6AB) especially in the beta range (12-30Hz); furthermore the change over time, i.e., the difference from the start of the block to the end of the block, was observable in the high-ASRS participants only (Figure 6CD). These results mirror our behavioral results from Experiment 1, where the low-ASRS group saw little difference in performance between music conditions and little change over time, whereas the high-ASRS group saw significant change over time across the music conditions (Figure 2B; significant interaction). Thus, in both brain and behavior, differences between Fast- and Control-Music were more apparent in high-ASRS individuals, and grew over the duration of the music; AM+Music both improved performance over time and stimulated neural activity in the beta range over time. This aligns with the idea that greater relative levels of beta-band activity are associated with improved performance on sustained attention tasks for individuals with ADHD^51–53^.

### Arousal does not explain effects of modulation across groups

While Experiment 1 suggested that the interaction between music for sustained attention and attentional deficit might depend on arousal, detailed examination of the effects of Fast- and Control-Music on phase-locking activity in the brain over time across ASRS groups (Figure 6CD) suggested that differences in acoustic modulation could underlie performance differences between high-arousal and low-arousal stimuli. In Experiment 4 the stimuli were made to vary only in modulation rate and depth, andhe condition with lowest arousal (16Hz medium depth condition) produced best performance in the high-ASRS group. Taken together the experiments suggest that acoustic modulation, rather than arousal per se, is the main driver behind the effects of music on sustained attention. Likewise, valence also cannot explain the differences in task performance. Although the increase in modulation depth from medium to high was sufficient to reverse the valence and arousal ratings of those stimuli, these ratings did not differ substantially between ASRS groups in the conditions that most affected sustained attention.

### Background music has widespread effects on cognitive control networks implicated in ADHD brain activity

fMRI results showed higher activity during AM+Music than during other acoustic conditions, with significant effects in widely distributed regions encompassing the salience, executive function, and default mode networks, especially in the high-ASRS group. When contrasting activity between hits and false alarms to isolate successful task-related activity, AM+Music elicited highest activity in the sensorimotor network, the salience network, and the visual association network. The widespread increases in activity during AM+Music confirm that background music affects sustained attention by influencing multiple interconnected networks that are normally coupled to subserve performance on a variety of cognitive tasks. The default mode and executive function network are typically anticorrelated in activity^54^, with the salience network being a consistent regulator of both other networks^29^. The interplay between salience, executive function, and default mode networks is critical in tasks requiring cognitive control, and this relationship is altered in ADHD by aberrant connectivity between these networks^55^. Here, when listening to AM+Music participants showed increased activity in the three networks. The interaction between group and music shows strong effects in the mid-cingulate cortex^56^ and the right anterior insula, key components of the salience network, especially in high-ASRS individuals during AM+Music. The simultaneously observed effects in the salience network and the late visual areas / ventral attention network may suggest that the AM+Music affects performance by motivating attention to the visual task. It may also suggest increased functional connectivity among the networks as neural activity becomes coupled to the music. Increased coupling may also explain results from the EEG experiment, which found increased phase-locking to the stimulus at specific frequencies that were targeted by the note rate and the added amplitude modulation patterns of the AM+Music. Within the phase-locking patterns across different frequencies during AM+Music, scalp topography showed highest phase locking around frontocentral channels at low frequencies (8Hz) but more widespread activity across the scalp at higher frequencies (32Hz). These topographical differences in phase-locked activity across different frequencies may be explained by cross-frequency coupling mechanisms which are known to underlie communication across different regions in the brain^57^.

### Oscillatory mechanisms mediate effects of music on sustained attention

Our subsequent behavioral results also point to the possibility that the music’s differential impact on oscillatory activity could underlie its effect on sustained attention. In Experiment 4 our stimuli were parameterized by modulation rate and showed a distinct 16Hz benefit for high-ASRS listeners, which could point to this effect being mediated by oscillatory processes in the brain. Individuals with ADHD have atypical oscillatory activity^36^, including a higher theta/beta ratio^58^. The observed rate-specific effect confined to the high-ASRS group in Experiment 4 could indicate that the AM+Music impacts the atypical brain activity in ADHD. Taken together our results suggest that there could be a process by which music at high modulation rates drives brain activity to benefit ADHD-like individuals in particular. Importantly, such modulation rates are not normally found in music, but were added into the AM+Music.

Brain stimulation methods with controlled frequencies have been used to enhance cognitive performance in recent years^59–62^. Sound is another means by which to stimulate the brain^63–65^; however, the frequency of sound stimulation has often resulted in stimuli that are perceptually unpleasant (e.g., click trains, binaural beats). The observation that music strongly affects neural oscillations^32,65,66^ motivates an approach whereby targeted acoustic modulation added to music might be used for neuromodulation, with particular goals outside the aesthetic and/or social uses of music (i.e., ‘functional’ music rather than ‘art’ music). One barrier to this has been individual variability, as different neurotypes and cognitive functions may require a range of targets for their mechanisms. Here we targeted subpopulations (by ASRS/ADHD) with hypotheses based on theoretical considerations from systems neuroscience, and found effects on brain and behavior that have implications for the use of music to enhance cognition in everyday life.

Fast and slow modulations in music can differently affect sustained attention. Results are not explained by valence and arousal, and may instead be explained by differences in phase-locking in specific frequencies as well as higher activity in widespread regions across the brain during task performance. Individuals who self-report ADHD symptoms are most sensitive to these modulations. This suggests that amplitude modulations in music could be used specifically to mitigate the negative effects of cognitive failures in sustained attention that come from attentional difficulties.

## Methods

### Experiment 1: Behavioral study

Experiment 1 presented three music background conditions to each participant (i.e., a within-subjects design), in randomized order. Each of the three conditions lasted 6 minutes and 54 seconds, the time required to complete 360 trials of the SART; the experiment ran 1080 trials in total (approximately 21 minutes).

#### Participants

We used Amazon’s Mechanical Turk platform to recruit and enroll participants. 114 participants were recruited for Experiment 1 (67 male, 40 female, 7 other/chose not to respond; mean age = 34). 102 participants were recruited to obtain valence and arousal ratings for the stimuli used in Experiments 1 and 4 (obtained in the same participants). Participants were asked to wear headphones, and not to change their volume or turn off audio during the experiment. They were told that the background music was unrelated to the task, but that they should nonetheless ensure they could hear the background music, because it was needed to control the acoustic environment across participants. To ensure compliance we employed a headphone screening task ^67^ as well as audio checks after each block, and a test of volume level at the end of the experiment. Participants who failed any of these were removed from data analysis. The final dataset for Experiment 1 comprised 87 of the initial 114 participants (76% passed screenings); the final dataset for the valence and arousal ratings comprised 62 of the initial 102 participants (60.7% passed screenings);

#### Stimuli

For Experiments 1 to 3, we chose as background auditory stimuli two commercially-available tracks of music that were predicted to span different arousal levels while being similar in valence levels, in addition to a Pink noise control stimulus. Background auditory stimuli were selected from commercially available options used to help people focus while working. The tracks we used were: (1) ‘Techno March’ by Brain.fm (high-arousal track) and (2) ‘Tracking Aeroplanes’ by The Echelon Effect (low-arousal track). Our listener ratings (Figure 1A) confirmed that the tracks differed in arousal but not in valence, suggesting that their effects on performance were attributable to arousal rather than valence. (3) Pink noise was also chosen as a control stimulus; it is often used for focused work while also being used in behavioral experiments because it has a spectrum that falls off with increasing frequency, similar to many auditory environments (unlike white noise which is spectrally flat). We limited our auditory stimuli to these three tracks in Experiments 1 in order to obtain experimentally well-validated auditory stimuli for neuroimaging and electrophysiology experiments in Experiments 2 and 3, both of which necessitated time-locked analyses on a smaller number of participants.

The tracks were musically dissimilar (to drive differences in valence and arousal) but in terms of low level acoustic features they differed most strikingly in the modulation domain--the less arousing music was at a slow tempo and contained slow modulations, while the more arousing music was at a fast tempo and contains fast modulations. The difference in the tracks’ modulation characteristics can be visualized with the modulation spectrum (Figure 1). The slow music contained a broad region of modulation energy from 0-8Hz, while the fast music contained a clear peak around 14-16Hz. This suggests tempo was not the sole cause of the difference in modulation characteristics, as tempo would simply shift the modulation spectrum. The difference in shape of the modulation spectrum instead reflects differences in the music, particularly note values (event durations). That is, the fast music was at a faster tempo, but also had faster note values. In particular, the modulation peak in the fast music was due to a consistent 32nd-note-rate amplitude modulation that had been deliberately applied as a feature of this kind of focus music. In contrast, the slow music contained long notes at several durations (e.g., whole, half) resulting in the absence of a single modulation peak. To complement the music tracks, the Pink Noise contained modulation energy broadly distributed in the higher modulation ranges (roughness). The modulation-domain differences were large and may partly underlie the music’s effects, but the tracks varied in other ways (tonality, instrumentation, etc.) which likely contributed to arousal. Due to the differences in frequency content of the three stimuli, rms-normalization was not appropriate (resulting in a large difference in perceived loudness across the stimuli); instead the stimuli were loudness-normalized by ear which produced rms values (average of left and right channels) of 0.059, 0.061, and 0.014 for AM+, Control-Music, and Pink Noise respectively.

#### Procedure

Users enrolled via Amazon’s Mechanical Turk platform, and provided informed consent as approved by IRB # 120180271 of New England IRB. To enroll, users must be over 18 and have normal hearing by self-report. If they chose to participate in our experiment, they were directed to a cover page with consent documentation and a simple description of the task, followed by a page with payment information. If they still chose to participate, they initiated a volume calibration task in which they heard a train of 1kHz tones at alternating levels, 10dB apart. They were told to set the volume on their computer so that only every other tone was audible, and told not to change their volume after this calibration step. To ensure compliance a short task after the main experiment required the participant to count the audible tones in a decrementing series of 1kHz tones (−5dB per step); those who counted fewer tones than expected were excluded from analysis. Following the initial volume calibration participants were directed to a headphone screening task^67^ composed of six 3AFC questions (<1 min). If they passed the headphone screening they were directed to the main task instructions and could begin when ready.

In Experiment 1 our participants completed 1080 trials of a sustained attention to response task (SART)^27,28,68^. In this task, a single digit ranging from 0 to 9 appeared on the screen for each trial. Each digit was presented for 250 ms followed by a 900 ms mask, resulting in a 1150 ms inter-trial interval. Participants’ task was to respond to any digit except for 0. Instructions for the task were as follows: “Numbers will appear on the screen. If you see 1-9, hit any key; if you see 0 do not hit any key.” Participants were paid at a rate of $0.01 per correct response and -$0.10m per commission error (misses were given no pay; $0.00), resulting in an average of ∼$12/hr for the overall task. Participants are not penalized in any way for leaving, but any who did were not granted the performance bonus (this was communicated at the outset), and were excluded from analysis as incomplete data. Participants were told the experiment would run for about 20 minutes, and they should try to complete the entire experiment.

Valence and arousal ratings for these stimuli were obtained in a separate group of 62 participants under the same screening procedures. In this rating task, a webpage displayed several audio player bars, each corresponding to a music track. Under each player were two sliders (100-pixel resolution) with the ends of the sliders labeled ‘Positive’ - ‘Negative’ and ‘Calming’ - ‘Stimulating’. Participants were given as much time as they liked to listen to the tracks and decide on the placement of the sliders, and were permitted to move between tracks to better judge them relative to one another. The stimuli for Experiment 1 and Experiment 4 were rated for valence and arousal together (by the same participants).

#### Data Analysis

Raw data from Amazon’s Mechanical Turk were exported to Matlab for analysis. The dependent variable was accuracy (d-prime), and independent variables were music condition and time (first half versus second half of each block). To test the effects of individual differences in attention difficulties, as quantified by the ASRS^69^, we did a median split on all subjects’ ASRS scores, yielding a high-ASRS group (i.e. those with more ADHD-like symptoms) and a low-ASRS group (those with less ADHD-like symptoms).

### Experiment 2: fMRI study

#### Participants

34 Wesleyan undergraduates (16 males, 18 females; mean age = 20.4, SD = 1.94) participated for course credit.

#### Stimuli and Procedure

The same three background music conditions from Experiment 1 (AM+Music, Control-Music, and Pink noise) were used in the fMRI study. During task fMRI, participants completed the SART while listening to AM+Music, Control-Music, and Pink noise in counterbalanced order. All experiment conditions were the same as Experiment 1 except inter-trial interval was 1425 ms, which was set to be equivalent to 3 TRs (as TR = 475 ms). Before each block the volume of the auditory stimulus was adjusted to a comfortable level (by communicating with the experimenter).

##### MRI Acquisition

High-resolution T1 and functional images were acquired in a 3T Siemens Skyra MRI scanner at the Olin Neuropsychiatry Research Center at the Institute of Living. The anatomical images were acquired using a T1-weighted, 3D, magnetization-prepared, rapid-acquisition, gradient echo (MPRAGE) volume acquisition with a voxel resolution of 0.8 × 0.8 × 0.8 mm^3^ (TR = 2.4 s, TE = 2.09 ms, flip angle = 8º, FOV = 256 mm). Task functional MRI was acquired as 1268 contiguous fast-TR echo planar imaging (EPI) functional volumes (TR = 475 ms; TE = 30 ms; flip angle = 90, 48 slices; FOV = 240 mm; acquisition voxel size = 3 × 3 × 3 mm^3^), resulting in a sequence that lasted approximately 10 minutes. Each background music condition was administered in a 10-minute sequence, resulting in approximately 30 minutes of scan time for each participant.

#### Data analysis

##### MRI Preprocessing

Task and structural MRI preprocessing was carried out using the Statistical Parametric Mapping 12 (SPM12) software (http://www.fil.ion.ucl.ac.uk/spm/) with the CONN Toolbox (http://www.nitrc.org/projects/conn)^71^. In order, this consisted of functional realignment and unwarp, functional centering, functional slice time correction, functional outlier detection using the artifact detection tool (http://www.nitrc.org/projects/artifact_detect), functional direct segmentation and normalization to MNI template, structural centering, structural segmentation and normalization to MNI template, and functional smoothing to an 8mm gaussian kernel ^72^. Denoising steps for functional connectivity analysis included correction for confounding effects of white matter and cerebrospinal fluid^73^, and bandpass filtering to 0.008-0.09 Hz.

##### Univariate Task-fMRI Analysis

Task fMRI analyses were done in SPM12 ^31^. Task fMRI analyses included: 1) Within-subject ANOVA comparing the three sessions (AM+, Control-Music, and Pink noise). 2) Hits, misses, false alarms, and correct rejections were separately modeled. 3) Seed-based functional connectivity was assessed using Conn and compared between the AM+, Control-Music, and Pink noise conditions.

##### Seed-Based Connectivity Analyses

Since we were interested in whole-brain connectivity patterns of known whole-brain networks, we used a functional network atlas that related 14 cortical networks to known cognitive functions^74^. Of these 14 networks, 7 aligned with our a priori hypotheses, including three DMN associated regions (vDMN, dDMN, Precuneus network), two ECN associated networks (LECN, RECN), and two salience associated networks (posterior salience network, anterior salience network). For the other 7 networks, hereafter referred to as post-hoc networks, we performed correction for multiple comparisons by dividing all p-value height thresholds by a factor of 7. Furthermore, noting a tendency towards false positives within resting state fMRI analyses ^75^, we used a conservative p-value height threshold of 0.001 for group comparisons and familywise error (p-FWE) correction for global effects of group and behavioral measures, so as to minimize the possibility of Type I Errors.

Mean time-course of each network was used as a single region-of-interest (ROI), and networks that resulted in clusters demonstrating a significant main effect of group were identified at the height threshold p < 0.05 FWE-corrected level. For each cluster that showed significant between-group differences, group-level beta values were extracted in order to determine the driving factors of these group differences. These networks were then used to extract group connectivity profiles at the height threshold p < 0.05, p-FWE corrected level.

### Experiment 3: EEG study

#### Participants

40 Wesleyan undergraduates (10 males; 30 females; mean age = 19, SD = 0.75) participated for course credit.

#### Stimuli and Procedure

Participants completed 680 trials of SART, a visual GO/NOGO task with inter-trial interval of 1150 ms, same as Experiment 1. This was done under four auditory conditions, presented in counterbalanced order: AM+, Silence (within-subjects), Control-Music, and Pink noise (between-subjects).

EEG was recorded with a 64-channel BrainVision actiCHamp system with PyCorder software in a sound attenuated and electrically shielded chamber.

#### Data analysis

Behavioral data: RT coefficient of variation (SD/M) was calculated for every block of 10 trials. EEG data were filtered using .5 Hz high pass filter and 60 Hz notch filter for electrical noise. Data were re-referenced to channels TP9 and TP10 and corrected for ocular artifacts using ICA. Matlab with EEGLAB toolbox ^76^ were used for analyses. Preprocessing: First, bad channels were rejected and then interpolated using EEGLAB’s rejchan and eeg_interp functions. Stimulus-brain coupling was assessed by first applying Morlet wavelet filtering at every single Hz from 1 to 50 Hz. Then, the Hilbert transform was done on each Hertz to get phase angle of stimulus and of EEG data, and the coupling between stimulus and EEG was assessed as the phase-locking value where plv(ch) = abs(sum(exp(1i*(phase_filtered_EEG(ch,:)- phase_filtered_music)))). This was applied in parallel to EEG data and to sound stimuli. The Pink Noise stimulus was generated in real time using Max/MSP (i.e., was not a stored file) and so the stimulus-brain coupling analysis was not applied to the Pink Noise data.

### Experiment 4: Behavioral study

#### Participants

Recruitment and experimental procedure (including screening) was identical to Experiment 1. Experiment 4 involved 221 participants (120 males; 99 females; 2 other/chose not to respond; mean age = 36, SD = 11.10). The final dataset for Experiment 4 comprised 175 of the initial 221 participants (79% passed screenings).

#### Stimuli

The stimuli (background music) were based on two different musical tracks; each had variants created that added amplitude modulation at three rates (8, 16, 32 Hz) and depths (low, medium, high). Modulation depth differences were quantified after processing to account for interactions between the music and modulator. We used the difference between original and processed tracks’ modulation spectra (in each cochlear channel; Fig 7) as a metric of applied modulation depth, and set the modulator such that our three depth conditions stepped up evenly in terms of this metric going from low to high depth. The transitions between conditions were implemented with smooth crossfades preserving track location (rather than a break and starting the music from the beginning).

Modulation patterns were aligned to the metrical grid of the music. This scheme meant that the relationship of the underlying music to the added modulation was as consistent as possible over very different rates. This is desirable in controlling for differences between conditions that arise from intrinsic properties of the underlying (pre-modulation) acoustic signal. For example, a musical event (e.g. a drum hit) transiently amplified by a modulation peak in the 8Hz condition would also be so in the higher-rate conditions. This is only the case because the different modulation rates are aligned to the music and are integer multiples of each other.

As can be seen in Figure 7, the applied modulation differences exist predominantly in the low-mid range of the frequency spectrum, with little modulation difference at high frequencies. This was by design, due to aesthetic considerations given the spectrotemporal density of the underlying music: Modulation applied in a broadband manner tended to interact with sparse events in the high frequency regions, which was occasionally annoying (salient). We confined added modulation to lower frequencies by applying it to only the frequency range 200Hz-1kHz.

Acoustic analysis before and after processing showed that the manipulated stimuli differed from the originals only in the modulation domain, and not in the audio frequency spectrum. Our conditions were therefore identical in terms of musical content and spectral balance (‘EQ’), eliminating important confounding factors and ensuring any behavioral differences can be attributed to applied modulation alone. Due to the similarity across the stimuli they were simply peak-normalized to 0.5, giving rms levels (average across left and right channels) in all cases between 0.080 and 0.092, where the faster and higher-depth modulation conditions had lower rms levels and the unmodulated tracks had the highest rms levels.

#### Procedure

Experimental procedure in Experiment 4 was similar to Experiment 1 with the addition of a fourth block of trials. The total number of trials in Experiment 4 was the same as in Experiment 1 (1080 trials) and so each block in Experiment 4 was 270 trials (∼5 minutes).

#### Data Analysis

For Experiment 4, a mixed-effects ANOVA was run on the dependent variable of accuracy, with the within-subjects factors of rate (4 levels: no-modulation, 8 Hz, 16 Hz, 32 Hz), and the between-subjects factors of ASRS (high vs. low) and extroversion (high vs. low).

## Supplementary Materials

### Listener volume settings

In Experiments 1 and 4, online participants were asked to first calibrate their volume with an alternating sequence of low-level tones separated by 10dB; they were told to set their computer volume so that only half the beeps could be heard. This approximate calibration was intended to somewhat narrow the distribution of volume settings participants used. In a post-test for volume, participants counted the audible tones in a series decrementing by 5dB per tone.

**Fig. S1.**
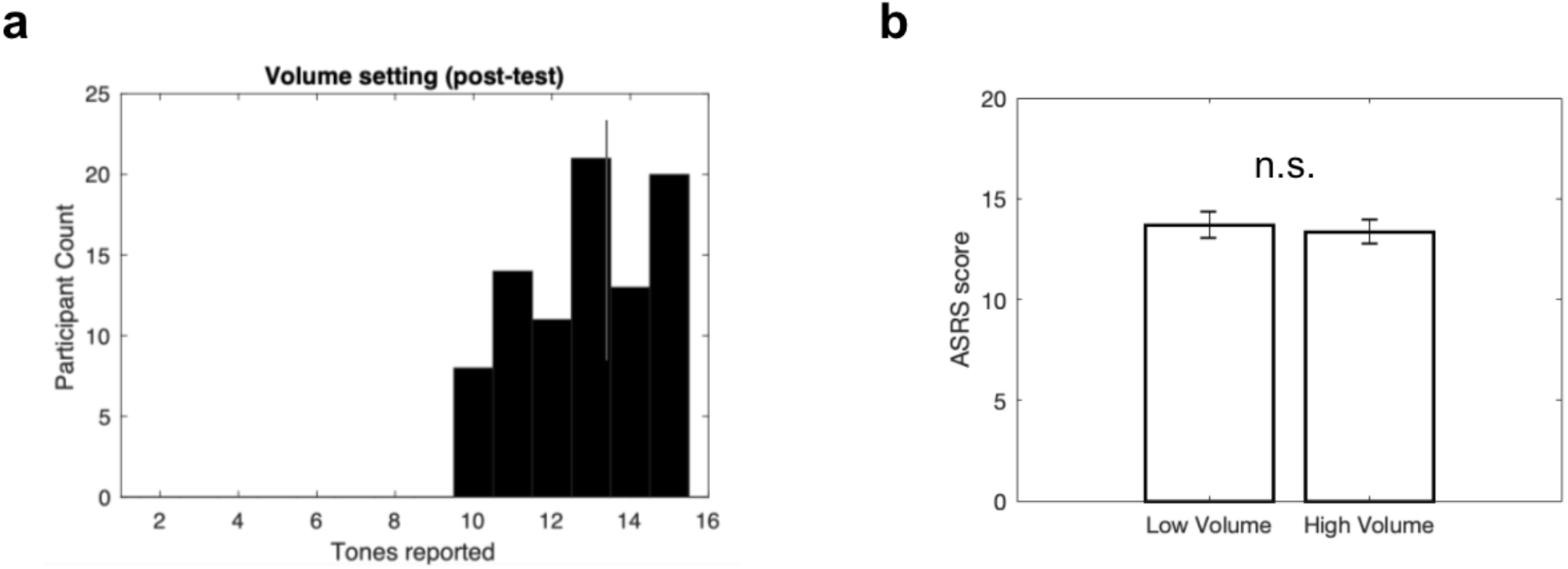
Online volume settings. **a**, Relative volume distribution for participants in Experiment 1 (N=87); measured as the distribution of tones counted in a volume-setting post-test where 15 tones decremented in volume by 5db per tone. **b**, ASRS scores across participants in the high- and low-volume groups.

Volume level might be expected to influence levels of arousal, or the effect of music in general ^78^. To see if volume levels interact with the factors addressed in our main results, we split our participants into two groups based on their volume settings (closest median split) and noted that the ASRS scores did not differ between the high-volume and low-volume listeners (Fig S1B).

**Fig. S2.**
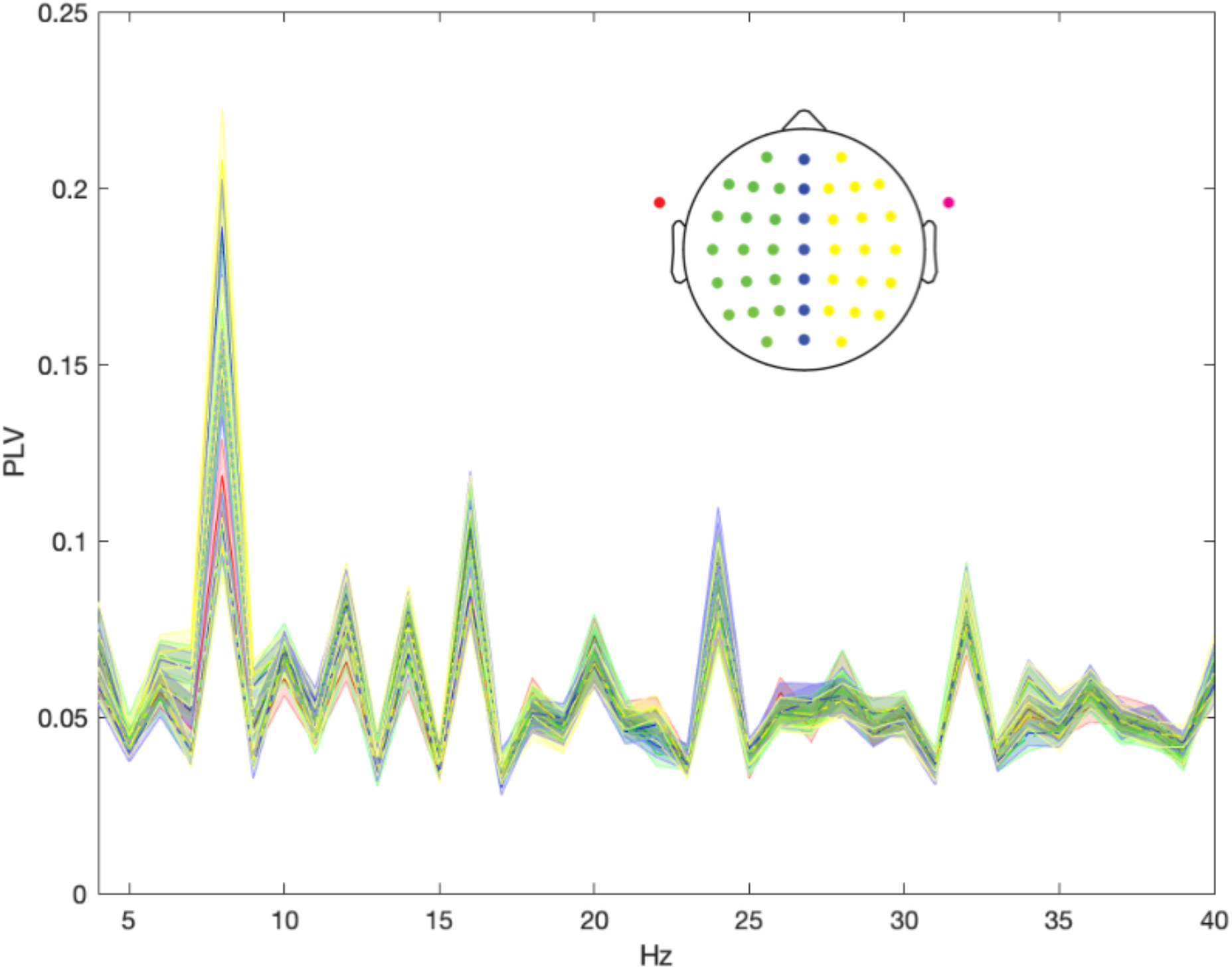
Stimulus-brain coupling across EEG electrode channels. For Experiment 3, PLV from EEG data are separately plotted here for left (green), midline (blue), and right (yellow) channels for frontal lateral sites (solid lines), frontal sites (dotted lines), and parietal sites (dashed lines). Results show largely similar stimulus-brain coupling across multiple recording sites.

